# Identification of active deubiquitinases in dendritic cells under unstimulated and activated conditions

**DOI:** 10.64898/2026.10.08.757166

**Authors:** Huw B Morgan, Jose A. Villadangos, Laura E. Edgington-Mitchell, Justine D. Mintern

## Abstract

Ubiquitination is a post-translational modification with well described roles in the regulation of immune cell function, such as dendritic cells (DCs). DCs are crucial initiators and regulators of T cell-mediated immune responses. In this study, we applied activity-based probes and proteomics to systematically profile DUBs in an immortalised DC line. Using the lysate-based biotin(bio)-Ubiquitin(Ub)-propargylamide(Prg) probe and proteomics, the activity and abundance of deubiquitinases were measured in unstimulated and CpG-stimulated MutuDCs to simulate responses to bacteria. This analysis was complemented with profiling using the cell-permeable DUB activity-based probe IMP-2373. The combined analysis with bio-Ub-Prg and IMP-2373 provided insights into the dynamics of DUB activity and abundance in DCs following stimulation, identifying several DUBs for further study.

## Introduction

Dendritic cells (DCs) are specialised antigen-presenting cells pivotal to adaptive immunity. These cells acquire and process antigens and are responsible for the initiation of T cell responses. Roles for ubiquitination machinery in regulating DC immunity are emerging to provide new insight into mechanisms of adaptive immunity. Ubiquitination is a post-translational modification involving the addition of ubiquitin (Ub) to lysine residues of proteins via a series of reactions involving E1-activating, E2-conjugating and E3 ligase enzymes (Komander et al. 2012)^1^. Ubiquitination is reversed by the actions of deubiquitinases (DUBs), which proteolytically cleave Ub from target proteins. There are approximately 100 known DUBs in humans, belonging to seven families: ubiquitin-specific peptidases (USPs), ubiquitin C-terminal hydrolases (UCH), Josephins (MJD), ovarian tumour proteases (OTU), MIU-containing DUBs (MINDY), zinc finger with UFM1-specific peptidase domain proteins (ZUP1) and Jad1/PAD/MPN domain-containing metalloenzymes (JAMM). Six of the seven families (USP, UCH, OUT, MJD, MINDY and ZUP1) are cysteine proteases and the seventh family (JAMM) are metalloenzymes (Clague et al. 2019)^2^. A small subset of DUBs have been identified as participating in DC immunity, including CYLD (Srokowski et al. 2009, Gringhuis et al. 2014, Wurm et al. 2015)^3–5^, TNFA1P3 (also known as A20, Kool et al. 2011, Liu et al. 2014, Xuan et al. 2015)^6–8^ and OTUB1 (Mulas et al. 2021)^9^ that are involved in NF-κB and MAPK signalling. Comprehensive profiling of DUBs that are active in DCs has not been previously undertaken. Currently, there is little information regarding the expression pattern of DUBs within DCs and how this is impacted after exposing DCs to inflammatory stimuli.

Here, we have used activity-based protein profiling (ABPP) to define the abundance and activity of DUBs in DCs. ABPP is a technique which uses an activity-based probe (ABP) to quantify enzymatic activity within lysates, cells or tissues. ABPs consist of three main elements: a reactive, electrophile often referred to as a ‘warhead’, a linker region and a reporter tag (Fang et al. 2021)^10^. The reactive warhead forms a covalent bond with the active site residue of an active enzyme. The cysteine protease DUBs can be targeted with covalent ABPs, including ubiquitin-vinyl sulfone (Ub-VS), which detected six of 17 known DUBs in yeast (Borodovsky et al. 2001)^11^. More advanced DUB ABPs have since been developed, including Ub-vinyl methylester (Ub-VME), which demonstrated broad DUB reactivity, and Ub-propargylamine (Ub-Prg), which exhibits enhanced specificity (Borodovsky et al. 2002, Ekkebus et al. 2013, Gorka et al. 2022)^12–14^. Due to their limited permeability, these probes must be used in cell lysates, and they have aided DUB profiling in cancers, identification of novel DUBs, structural and mechanistic studies and the screening and characterisation of inhibitors (Gorka et al. 2022, Pinto-Fernandez et al. 2019, Mahajan et al. 2022, Hewings et al. 2018, Kwasna et al. 2018, Grethe et al. 2022, Hermanns et al. 2022, Altun et al. 2011, Cookson et al. 2023, Hewings et al. 2017)^14–23^. Here we have used ABPP to define DUB expression and activity profiles in unstimulated and activated DCs.

## Results

### DUB expression profiles in unstimulated and activated dendritic cells

Analysis of DUBs in dendritic cells was assessed using the Mutu dendritic cell line (Marraco et al. 2012)^24^. First, we undertook an unbiased global cell proteomics approach to determine the range and abundance of DUBs expressed by unstimulated MutuDCs compared to those stimulated with 0.5 μM CpG for 18 hours at 37°C. Several DUBs exhibited differential abundance following activation (**Figure 1**, **Table 1**).

**Figure 1:**
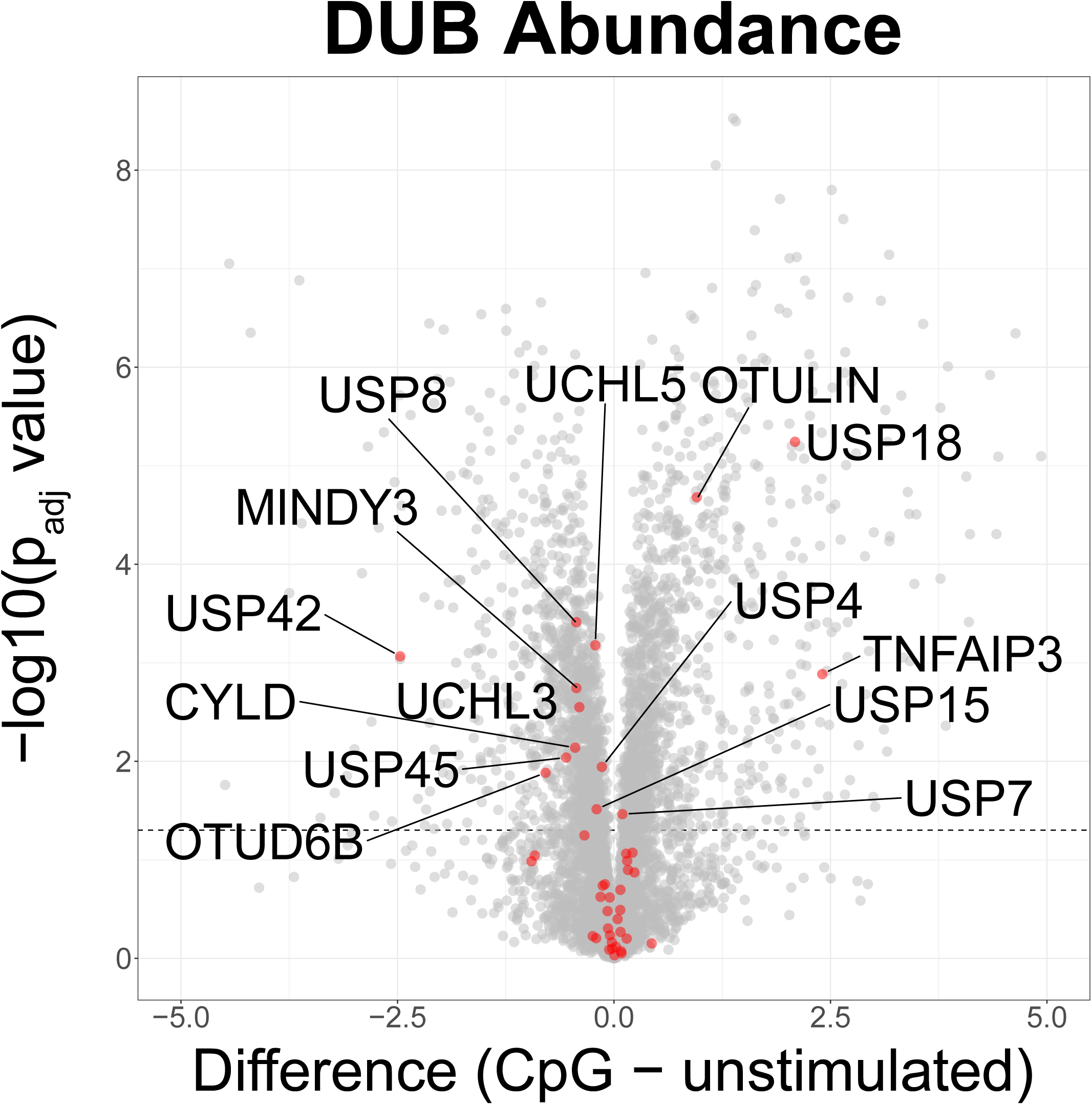
Stimulation with CpG alters the abundance of deubiquitinases in MutuDCs. MutuDCs were stimulated with 0.5 μM CpG for 18 hours at 37⁰C before lysis and tryptic digest for analysis by LC-MS/MS using label-free quantification. Volcano plot depicts the difference between CpG-stimulated and unstimulated MutuDCs. Data from four independent experiments, two-tailed Student’s T test with permutation-based false discovery rate with a significance threshold of padj > 0.05 (-log_10_ (p_adj_ value) > 1.3).

**Table 1:** Changes in deubiquitinase abundance and activity following MutuDC CpG stimulation relative to unstimulated cells.

| Deubiquitinase | Abundance |  | Activity |  |
| --- | --- | --- | --- | --- |
|  | Log2(Fold Change LFQ CpG:Unstimulated) | Log10(adj. P-value) | Log2(Fold Change LFQ CpG:Unstimulated) | Log10(adj. P-value) |
| OTUD1 | ND | ND | 1.516938 | 2.466088 |
| USP16 | 0.236320496 | 0.872502464 | 0.274388 | 1.459873 |
| OTUB2 | -0.952121258 | 0.985511245 | 0.225405 | 0.311029 |
| USP14 | -0.045850754 | 0.234253523 | 0.211416 | 0.460672 |
| USP42 | -2.469167233 | 3.065760001 | 0.117726 | 0.233454 |
| OTUB1 | -0.129567146 | 0.737667729 | 0.092297 | 0.07678 |
| USP30 | -0.245466709 | 0.225936462 | 0.082886 | 0.14921 |
| USP7 | 0.097499371 | 1.464147848 | 0.074178 | 0.312297 |
| USP20 | ND | ND | 0.051474 | 0.092813 |
| OTUD7B | 0.075712681 | 0.268210857 | 0.043755 | 0.170306 |
| USP38 | -0.068763733 | 0.303440954 | 0.025908 | 0.068223 |
| USP3 | 0.436130047 | 0.151115891 | 0.025373 | 0.052312 |
| USP40 | ND | ND | -0.01393 | 0.050204 |
| VCPIP1 | 0.074430943 | 0.695777655 | -0.01894 | 0.06701 |
| USP47 | 0.152627945 | 0.989848187 | -0.04917 | 0.168406 |
| UCHL5 | -0.215732098 | 3.17820508 | -0.06143 | 0.107541 |
| BAP1 | ND | ND | -0.07322 | 0.140423 |
| USP25 | 0.005135059 | 0.031383934 | -0.09352 | 0.194259 |
| USP22 | ND | ND | -0.11287 | 0.44707 |
| USP48 | 0.14700079 | 0.200894135 | -0.11798 | 0.266641 |
| USP4 | -0.139659405 | 1.943710873 | -0.14637 | 0.418932 |
| USP32 | 0.081841469 | 0.072267901 | -0.20109 | 0.381389 |
| YOD1 | ND | ND | -0.22094 | 0.407817 |
| OTUD5 | 0.161913395 | 0.897577307 | -0.25398 | 0.148635 |
| JOSD2 | ND | ND | -0.25454 | 0.310141 |
| USP9X | -0.024907112 | 0.163686658 | -0.25796 | 1.14376 |
| USP33 | -0.206184864 | 0.204773621 | -0.2594 | 0.981694 |
| USP36 | ND | ND | -0.28122 | 0.948235 |
| USP15 | -0.200268269 | 1.513811695 | -0.28435 | 1.186319 |
| USP19 | 0.072498322 | 0.491267735 | -0.28724 | 0.699057 |
| OTUD6B | -0.789832115 | 1.88414973 | -0.29955 | 0.696193 |
| OTUD4 | 0.02200985 | 0.117416375 | -0.30259 | 1.123234 |
| USP28 | ND | ND | -0.30398 | 0.412189 |
| USP5 | -0.049475193 | 0.618126485 | -0.33792 | 0.976741 |
| UCHL3 | -0.399524212 | 2.549658371 | -0.35641 | 0.174292 |
| USP45 | -0.552838802 | 2.037892753 | -0.39141 | 1.701179 |
| USP12 | -0.91397953 | 1.044041765 | -0.46168 | 0.784645 |
| USP10 | -0.104595184 | 0.752331055 | -0.47801 | 1.999595 |
| USP46 | ND | ND | -0.48113 | 0.58078 |
| USP8 | -0.435637474 | 3.412753774 | -0.53225 | 3.480466 |
| USP24 | -0.154959202 | 0.624111493 | -0.53814 | 2.65426 |
| USP1 | ND | ND | -0.61037 | 3.204167 |
| USP34 | -0.054656982 | 0.088441189 | -0.61279 | 1.10336 |
| CYLD | -0.448182583 | 2.138420455 | -0.62701 | 2.151525 |
| USP21 | ND | ND | -0.74211 | 0.978721 |
| USP37 | 0.087571621 | 0.052220779 | -1.35136 | 4.125889 |
| TNFAIP3 | 2.405446053 | 2.884987562 | ND | ND |
| USP18 | 2.090750694 | 5.24350655 | ND | ND |
| OTULIN | 0.956209183 | 4.679959294 | ND | ND |
| USP39 | 0.213591576 | 1.072714147 | ND | ND |
| OTULINL | 0.140624046 | 1.062276191 | ND | ND |
| PSMD14 | 0.042272568 | 0.39876605 | ND | ND |
| EIF3F | -0.02035284 | 0.098241228 | ND | ND |
| EIF3H | -0.074324131 | 0.479922933 | ND | ND |
| ATXN3 | -0.340889454 | 1.24822721 | ND | ND |
| MINDY3 | -0.434487343 | 2.743240373 | ND | ND |

Green = upregulated, $\log_{10}(\text{adj. P-value}) > 1.3$ ; red = downregulated, $\log_{10}(\text{adj. P-value}) > 1.3$ ; ND = not detected.

### Identifying active DUBs in unstimulated and activated dendritic cells using a cell-impermeable activity-based probe

Next, we used the ABP biotin-ubiquitin-propargylamide (Bio-Ub-Prg) to profile DUB activity in naïve and activated MutuDCs. To confirm the selectivity of the biotin-ubiquitin (Ub)-propargylamide probe for active DUBs, we compared Bio-Ub-Prg-treated lysates of naïve and CpG-stimulated MutuDCs in the presence and absence of N-ethylmaleimide (NEM), a broad spectrum cysteine protease inhibitor. Probe-labelled proteins were affinity purified with streptavidin-agarose beads and subjected to trypsin digestion and LC-MS/MS. We observed loss of DUB detection in NEM samples, confirming selectivity for active enzymes as expected (**Supplementary Figure 1**).

Bio-Ub-Prg was used to profile changes in DUB activity following MutuDC activation. We first undertook analysis by SDS-PAGE and western blotting to assess potential changes in DUB activity in MutuDCs exposed to an array of different stimuli. MutuDCs were examined at rest or following activation by different stimuli, including 0.5 μM CpG, 100 μg/mL poly I:C or 50 ng/mL IFNγ for 18 hours followed by lysate labeling with Bio-Ub-Prg, with or without NEM pre-treatment. ABP-treated cell lysates were subjected to SDS-PAGE and streptavidin blotting. A distinct pattern of probe-labelled proteins of varying molecular weight was observed in lysates generated from unstimulated or activated MutuDCs, with subtle differences detected across stimuli (**Figure 2**).

**Figure 2:**
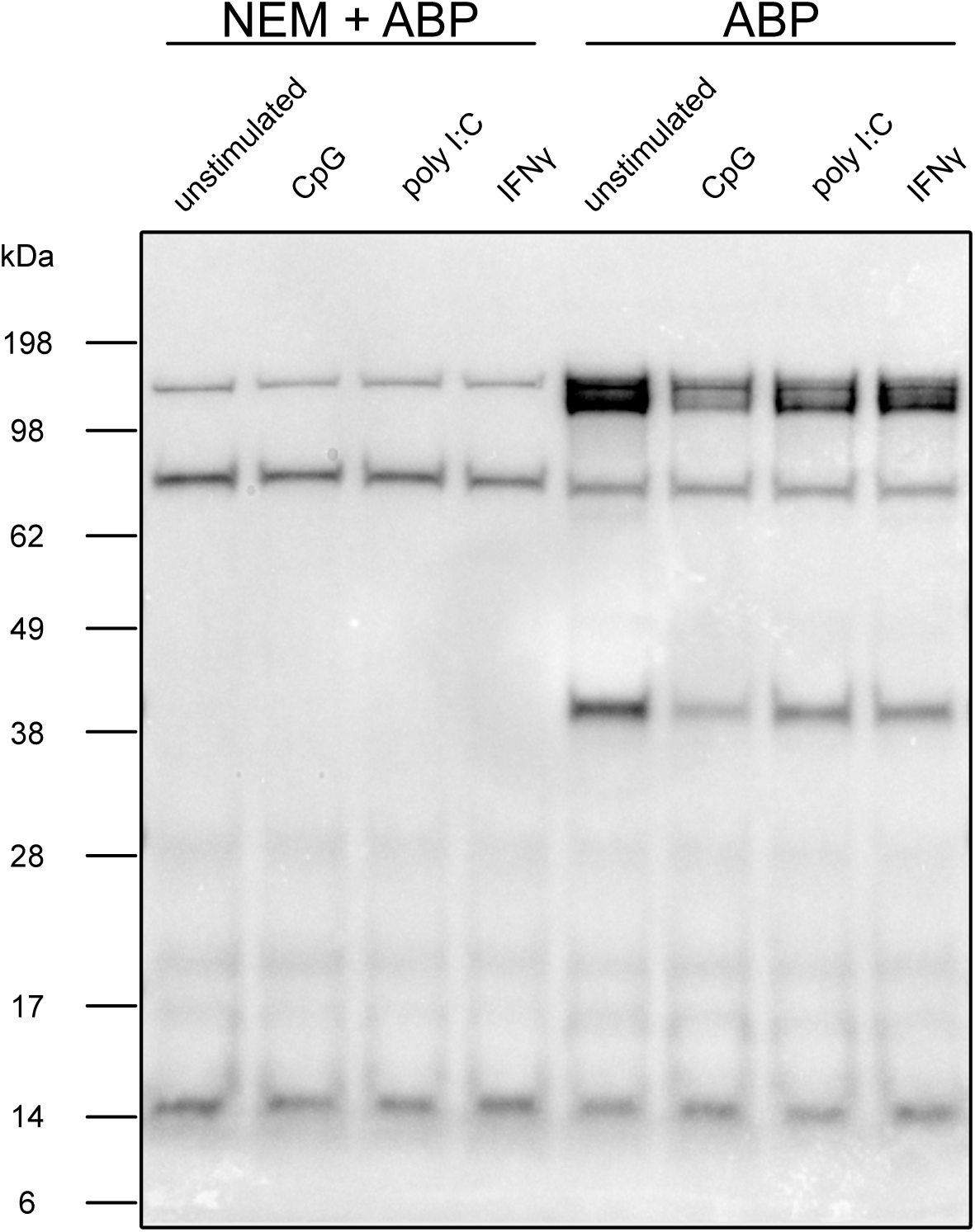
MutuDCs display altered deubiquitinase activity following inflammatory stimulation. MutuDCs were stimulated with 0.5 μM CpG, 100 μg/mL poly I:C, 50 ng/mL IFNγ, or 0.25 μM CpG and 50 ng/mL IFNγ for 18 hours at 37⁰C. Cells were lysed and protein was quantified by bichinoic acid assay. 50 μg of lysate protein was incubated with 0.1 μg Bio-Ub-Prg activity-based probe (ABP) for 1 hours at 37⁰C. Where indicated, samples were pre-incubated with N-ethylmaleimide (NEM) for 1 hours prior to ABP addition. Equal amounts of protein were loaded for analysis of probe binding by SDS-PAGE. Blot shown is representative of three independent experiments.

To gain detailed insight into how DUB abundance and activity varies in unstimulated versus stimulated MutuDC, we selected CpG-activated conditions for further analysis. LC-MS/MS was performed on whole cell lysates to measure DUB abundance and following affinity purification of Bio-Ub-Prg ABP-treated lysates to measure DUB activity. Comparison of DUB activity and abundance between unstimulated and CpG-stimulated MutuDCs yielded a list of differentially active and abundant DUBs. Thirty-four DUBs had detectable protein in both the pulldown and global proteomics datasets. Generally, the activity of the DUBs correlated with the abundance of the DUB (**Figure 3A**). Several DUBs exhibited differential activity between unstimulated and stimulated cells (**Figure 3B**, **Table 1**) which correlated with abundance such as CYLD and USP8, while many were altered only at the activity level, such as USP16, USP24, USP37 and USP45 (**Figure 3B, C, Table 1**). Some DUBs (USP1 and OTUD1) had detectable activity but were not detected in the global proteomics dataset, potentially due to their low expression levels (**Figure 3B, C**). Other DUBs, such as OTULIN, TNFAIP3, USP18 and USP42, exhibited significantly altered abundance following activation but were not labelled by the ABP **(Figure 3B, C).** Some of these DUBs are known to have non-ubiquitin substrates (USP18) or have poor reactivity with monoubiquitin ABPs (OTULIN, TNFAIP3) (Hewings et al. 2018, Honke et al. 2016, Zhang et al. 2017, Mevissen et al. 2013)^17,25–27^.

**Figure 3:**
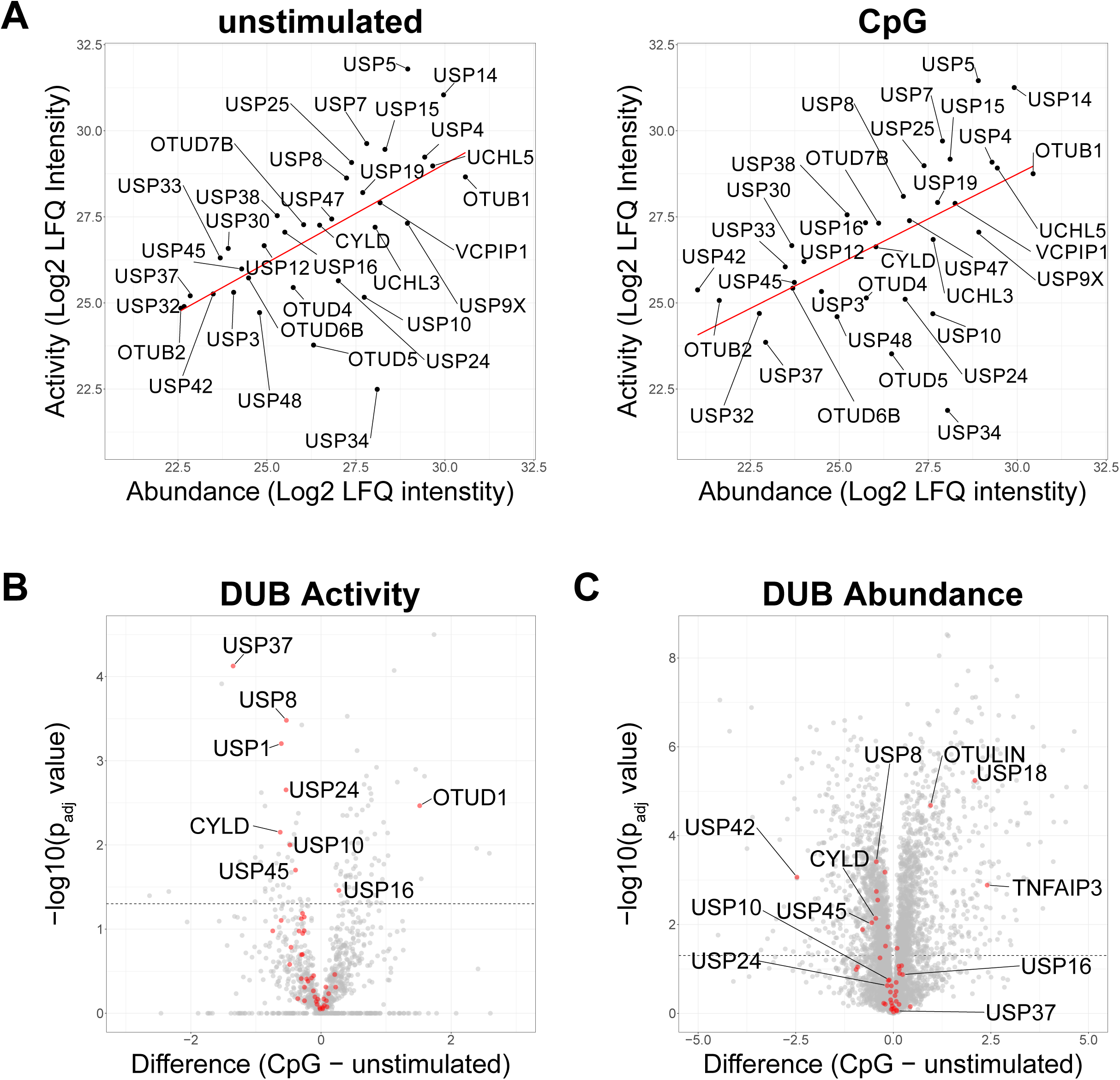
Activity-based protein profiling with Bio-Ub-Prg reveals changes in deubiquitinase activity in MutuDCs independent of deubiquitinase abundance. MutuDCs were stimulated with 0.5 μM CpG for 18 hours at 37⁰C. Cells were lysed and lysate protein was quantified by bichinoic acid assay. 500 μg of lysate protein was incubated with 0.1 μg Bio-Ub-Prg activity-based probe (ABP) for 1 hours at 37⁰C. 20 μg of lysate protein was set aside for global proteome analysis. Streptavidin-agarose beads were used to pulldown probe-labelled proteins. Streptavidin-agarose beads were subjected to on-bead trypsin digest prior to analysis by mass spectrometry. A) Scatter plots of deubiquitinase (DUB) activity and DUB abundance for proteins with detected protein in activity and abundance proteomics. Red line indicates correlation between activity and abundance. Volcano plots of B) DUB activity and C) abundance. Data from four independent experiments, two-tailed Student’s T test with permutation-based false discovery rate with a significance threshold of p_adj_>0.05 (-log_10_(p_adj_ value)>1.3).

### Identifying active DUBs in unstimulated and activated dendritic cells using a cell permeable activity-based probe

In addition to being incapable of detecting DUBs with non-ubiquitin substrates, Bio-Ub-Prg requires cell lysis disruption of physiological environment prior to labelling, potentially altering their activity (Hewings et al. 2017)^23^. To address these issues, a cell-permeable ABP called IMP-2373 was used to profile DUBs in MutuDCs. IMP-2373 consists of a pyrrole benzylamide backbone with a cyanopyrrolidine warhead, which covalently reacts with the catalytic cysteine, and a terminal alkyne for addition of biotin by copper-catalysed azide-alkyne cycloaddition (CuAAC) click chemistry (Conole et al. 2023)^28^. Cells were treated with 10 μM IMP-2373 or DMSO vehicle control for 1 hour before lysis and biotin addition by click chemistry. As DUB labelling was performed prior to lysis, a detergent-based lysis method was used. IMP-273-labelled proteins were then affinity purified with streptavidin-agarose beads. Successful labelling was confirmed by streptavidin blot. Bands in DMSO treated cells were likely to be endogenously biotinylated proteins or those non-specifically labelled by click chemistry (**Supplementary Figure 2**). We then applied IMP-2373 to profile DUB activity in unstimulated and CpG-stimulated MutuDCs using LC-MS/MS. DMSO-treated cells were used as a control for non-specific biotin labelling and streptavidin-agarose interactions. MutuDCs were kept at rest or stimulated with 0.5 μM CpG for 17 hours followed by incubation with 10 μM IMP-2373 or 0.01% DMSO vehicle control for 1 hour. After biotin addition, proteins were subjected to on-bead digest and LC-MS/MS. A pre-enrichment aliquot of lysate protein was analysed in parallel to assess total DUB abundance. As IMP-2373 is a small molecule, it has potential to be cytotoxic. To determine if IMP-2373 induced cytotoxicity and altered the cellular profile, we compared the global proteome of IMP-2373 and DMSO-treated MutuDCs. There were no significant changes in the proteome between IMP-2373 and DMSO-treated MutuDC when unstimulated or following CpG stimulation indicating that any changes in DUB activity or abundance detected in subsequent analyses was not due probe-mediated cellular responses or cytotoxicity (**Figure 4A**). Comparison of proteins biotinylated proteins from IMP-2373 and DMSO-treated cells also indicated that there was little non-specific pulldown of DUBs and that purified DUBs pulled were labelled with IMP-2373 (**Figure 4B**).

**Figure 4:**
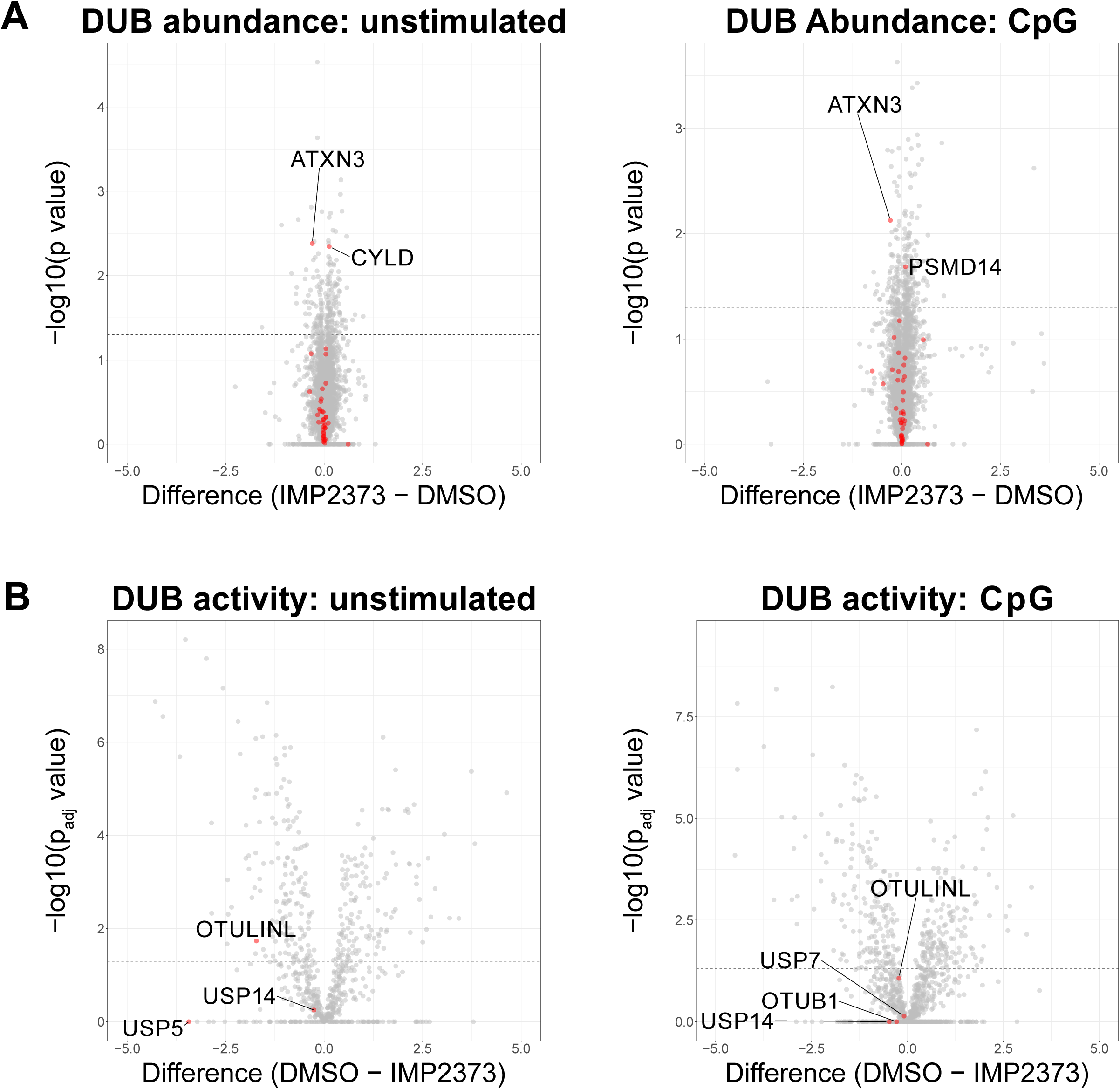
IMP-2373 does not alter the proteome of MutuDCs and deubiquitinases are not pulled down in the absence of IMP-2373. MutuDCs were stimulated with 0.5 μM CpG for 17 hours at 37⁰C. Cells were treated with DMSO or 10 μM IMP-2373 for 1 hour at 37⁰C. Cells were lysed and lysate protein was quantified by bichinoic acid assay. 500 μg of lysate protein from DMSO- and IMP-2373-treated cells were then subject to CuAAC click chemistry addition of biotin to IMP-2373 labelled-proteins. 20 μg of lysate protein was set aside for global proteome analysis. Streptavidin-agarose beads were used to pulldown biotin-labelled proteins. Streptavidin-agarose beads were subjected to on-bead trypsin digest prior to analysis by mass spectrometry. A) Volcano plots of A) DUB abundance and B) activity. Data from five independent experiments, two-tailed Student’s T test with permutation-based false discovery rate with a significance threshold of p_adj_>0.05 (-log_10_(p_adj_ value)>1.3).

Comparison of DUB activity and abundance between unstimulated and CpG-stimulated MutuDCs using IMP-2373 was undertaken. DUBs were detected in the immunoprecipitation and global proteomics for unstimulated and CpG-stimulated MutuDC, (25 and 26 respectively). Unlike with Bio-Ub-Prg, DUB activity did not correlate with abundance in unstimulated or CpG-stimulated MutuDCs (**Figure 5A**). ATXN3 was more abundant in unstimulated MutuDC, while TNFAIP3 and USP18 were significantly increased in abundance following CpG stimulation, and COPS5 displayed no significant alternation (**Figure 5B**). Both ATXN3 and COPS5 had elevated activity in CpG-stimulated MutuDCs, with USP18 identified as significantly more active in unstimulated cells (**Figure 5C**).

**Figure 5:**
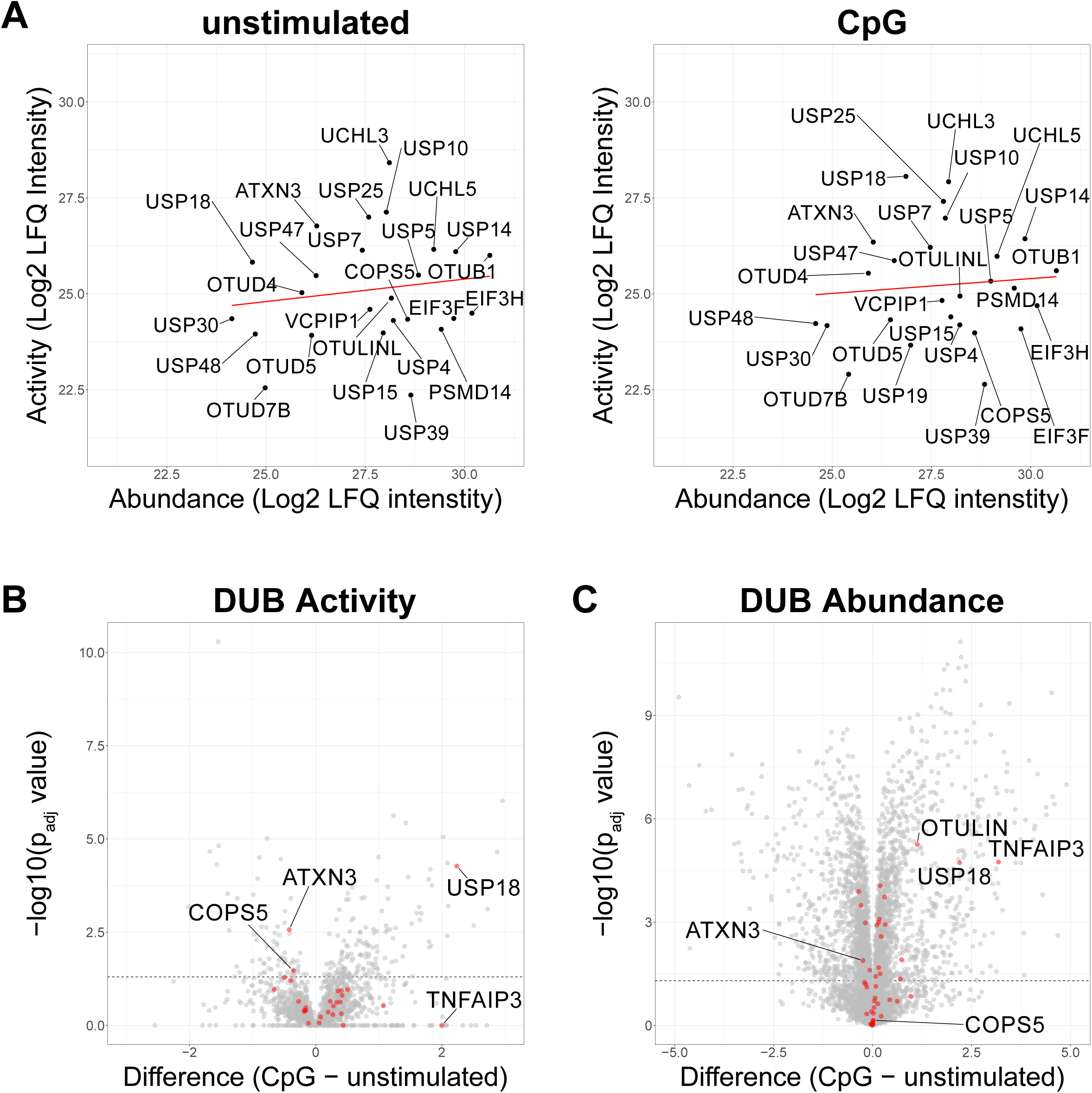
Activity-based protein profiling with IMP-2373 detects changes in DUB activity in MutuDCs not captured by Bio-Ub-Prg. MutuDCs were stimulated with 0.5 μM CpG for 17 hours at 37⁰C. Cells were treated with DMSO or 10 μM IMP-2373 for 1 hour at 37⁰C. Cells were lysed and lysate protein was quantified by bichinoic acid assay. 500 μg of lysate protein from DMSO- and IMP-2373-treated cells were then subject to CuAAC click chemistry addition of biotin to IMP-2373 labelled-proteins. 20 μg of lysate protein was set aside for global proteome analysis. Streptavidin-agarose beads were used to pulldown biotin-labelled proteins. Streptavidin-agarose beads were subjected to on-bead trypsin digest prior to analysis by mass spectrometry. A) Scatter plots of deubiquitinase (DUB) activity and DUB abundance for proteins with detected protein in activity and abundance proteomics. Red line indicates correlation between activity and abundance. Volcano plots of B) DUB activity and C) abundance. Data from four independent experiments, two-tailed Student’s T test with permutation-based false discovery rate with a significance threshold of p_adj_>0.05 (-log_10_(p_adj_ value)>1.3).

## Discussion

Here, we have combined global proteomics and analysis using ABPP to generate detailed profiles of DUBs in unstimulated and activated DCs. We have gained in-depth insight into DUB abundance and activity in unstimulated cells and observed how they are impacted following DC stimulation.

In this study we have used two distinct ABPs to conduct our analysis; Bio-Ub-Prg and IMP-2373. The differences in generated outcomes likely stem from Ub being the targeting moiety for Bio-Ub-Prg, compared to the pyrrole benzylamide backbone of IMP-2373. The Ub moiety restricts Bio-Ub-Prg to use within cell lysates, potentially altering DUB activity outside of their physiological context. Moreover, use of Ub restricts Bio-Ub-Prg to detection of Ub-specific DUBs and excludes those specific for Ub-like modifications including ISGylation, neddylation and SUMOylation. This is highlighted in our analysis where COPS5 and USP18, a deneddylating and deISGylating enzyme respectively, were only detected using IMP-2373. Overall, when both datasets were considered, 13 differentially active DUBs were identified comparing unstimulated and CpG-stimulated MutuDCs. In unstimulated DCs, active DUBs were ATXN3, COPS5, CYLD, USP1, USP8, USP10, USP24, USP37 and USP45. Several of these, ATXN3, USP1, USP8, USP10, USP37 and USP45, are involved in regulating DNA damage and cell cycle progression, thus their elevated activity in unstimulated MutuDCs is expected as TLR stimulation suppresses MutuDC proliferation (Fuertes Marraco et al. 2012, Garcia-Santisteban et al. 2013, Hernandez-Carralero et al. 2024, Song et al. 2025, Bhattacharya et al. 2020, Chauhan et al. 2021, Li et al. 2023)^24,29–34^. CYLD is a DUB with well-known roles in DC immunity (Srokowski et al. 2009, Wurm et al. 2015, Gringhuis et al. 2014)^3–5^. Our profiling fits with the known activity pattern of CYLD in DCs, where following DC activation, the full length CYLD is replaced by a shorter isoform (sCYLD) that cannot bind its allosteric activator (Elliott et al 2021)^35^. Another well-characterised DUB in DC function and a negative regulator of NF-κB signaling is TNFAIP3. TNFAIP3 was both more abundant and more active in the CpG-stimulated MutuDCs. Upregulation of TNFAIP3 following TLR4 stimulation of DCs has been previously reported (Liu et al. 2014)^7^. TNFAIP3 and CYLD share similar substrates in DCs, namely core components of the NF-κB signaling pathway NEMO, RIPK1 and TRAF6. USP16, USP18 and OTUD1 demonstrated elevated activity following CpG stimulation of MuTuDCs. USP16 is as a positive regulator of NF-κB signalling (Yu et al. 2016)^36^, while OTUD1 has mixed roles as a negative or positive regulator of inflammatory NF-κB signalling (Oikawa et al. 2022, Wu et al. 2022, Zhu et al. 2025)^37–39^. Whether USP16 and OTUD1 play a similar role in regulating NF-κB signaling in DCs would need to be confirmed with knockout studies. Several DUBs with known roles in DC function were not captured by our activity-based probe profiling with either Bio-Ub-Prg or IMP-2373. These include YOD1, MYSM1, USP21, ZRANB1 and UCHL1 and this may be a consequence of the use of the MutuDC cell line. OTUD7B was detected in both the global proteomics and with both ABPs but there was no differential OTUD7B activity or abundance detected between unstimulated and CpG-stimulated MutuDC correlating with published data (Harit et al. 2023)^40^.

In the context of immune cells, very few studies have undertaken ABPP of DUBs. In T cells, HA-Ub-VME was used to profile DUB activity downstream of T cell receptor (TCR) engagement (Jahan et al. 2016)^41^. These experiments identified increased activation of USP12 following TCR engagement and concomitant with its migration from the nucleus to the cytosol where it associated with the TCR. DUB ABPs have also been used to study changes in DUB activity upon exposure and infection of macrophages with bacterial pathogens such as *Salmonella typhimurium*, *Yersinia enterocilica* and *Francisella tularensis* (Kummari et al. 2015, Santelices et al. 2023, Vozandychova et al. 2023)^42–44^. ABPP identified reduced activity of USP8 and UCHL5 following *Salmonella* infection. Loss of USP8 activity promoted bacterial survival by downregulating autophagy, while loss of UCHL5 activity leads to reduced IL-1β release following inflammasome activation (Kummari et al 2015, Santelices et al. 2023)^42,43^. In *Francisella*-infected macrophages, the activity of USP10, USP25 and UCHL5 was reduced but protein levels were stable relative to uninfected cells (Vozandychova et al. 2023)^44^.

In summary, we have conducted detailed profiling of DUB abundance and activity in unstimulated and activated DCs. This opens potential avenues for further investigation of the role of DUBs in regulating DC function.

## Methods

### Cell culture

MutuDC1940 cells (MutuDCs) were a gift from Hans Acha-Orbea (Marraco et al. 2012)^24^. Cells were maintained in IMDM supplemented with 10% FBS, 50 μM β-mercaptoethanol and 100 U/ml penicillin and 100 μg/ml streptomycin, and cultured at 37°C, 10% CO_2_. Once cells reached 70% confluency, cells were subcultured using EDTA-BSS (balanced salt solution) supplemented with 2% FCS. Adherent and non-adherent cell populations were collected and pooled for subculturing.

### Activity-based Profiling of DUBs with Bio-Ub-Prg

30 x 10^6^ MutuDCs were seeded in two 175-cm^2^ flasks. One flask was stimulated with 0.5 μM CpG for 18 hours at 37⁰C, 10% CO_2_. MutuDCs were harvested using EDTA-BSS 2% FCS and washed three times with PBS. Cells were resuspended in sonication buffer (50 mM Tris-HCl pH 7.4, 5 mM MgCl_2_, 0.5 mM EDTA, 250 mM sucrose, 1 mM dithiothreitol) and lysed by sonication (8 cycles, 5s on, 30s off, 35% power). Lysate was cleared and protein quantified using a Pierce BCA assay kit (ThermoFisher). Total protein (500 μg) was aliquoted into two samples and a minimum of 20 μg was analysed by global proteomics. One 500 μg aliquot was treated with 20 mM N-Ethylmaleimide (NEM), a broad-spectrum cysteine protease inhibitor, and both were incubated for 1 hour at 37⁰C. Bio-Ub-Prg (1 μg; UbiQ) was added to each sample. Labelling was quenched by addition of 0.4% SDS and 0.5% NP-40 substitute. Pierce high-capacity streptavidin agarose resin (20 μL slurry: ThermoFisher) was added to each sample and incubated for 1.5 hours at 4⁰C with end-over-end agitation. Beads were washed four times with PBS. For streptavidin blot, beads were boiled in SDS sample buffer at 95⁰C for 5 minutes to elute and denature samples. For mass spectrometry, beads were washed further with 20 mM TEAB pH 8 before on-bead trypsin digestion.

### Activity-based Profiling of DUBs with IMP-2373

8 x10^6^ MutuDCs were plated in 10-cm^2^ plates. Two plates were stimulated with 0.5 μM CpG for 18 hours at 37⁰C, 10% CO_2_. One plate from each stimulation group was then treated with 10 μM IMP-2373 or the equivalent volume of DMSO for 1 hour at 37⁰C, 10% CO_2_. MutuDCs were harvested using EDTA-BSS 2% FCS and washed with PBS. Cells were lysed using IMP-2373 lysis buffer (1% [v/v] Triton-X, 0.1% [w/v] SDS, 1x cOmplete protease inhibitor cocktail [Roche] in PBS) for 20 minutes on ice. Lysed samples were vortexed and lysates cleared. Lysate protein was quantified using a Pierce BCA assay kit (ThermoFisher). Total protein (500 μg) was aliquoted for click chemistry labelling and 100 μg analysed by global proteomics. For click chemistry, each sample was adjusted to 0.5-2 mg/mL protein concentration with PBS to a volume of 300 μL. A click mixture of 3 μL biotin-azide (10 mM stock), 6 μL copper sulphate (50 mM stock), 6 μL TCEP (50 mM stock) and 3 μL TBTA (10 mM stock) was prepared and incubated for two minutes at room temperature. The click mixture was added to each sample and incubated for 1 hour on a shaker at room temperature. The click reaction was quenched using 10 mM EDTA and residual click reagents removed from the sample using methanol-chloroform precipitation. Samples were then prepared for streptavidin blotting or affinity enriched and prepared for LC-MS/MS as previously described

### On-bead trypsin digest and liquid chromatography tandem mass spectrometry

Proteins were reduced on-bead with 5 mM TCEP in 20 mM TEAB for 30 minutes at 45⁰C. Iodoacetamide was added to a final concentration of 20 mM. Samples were incubated for 10 minutes at room temperature in the dark. Following reduction and alkylation, beads were spun down and supernatant removed. Beads were resuspended in 200 μL 20 mM TEAB containing 500 ng trypsin protease (ThermoFisher) and incubated for 15 hours at 37⁰C on a shaking incubator. Following digestion, samples were spun down and supernatant harvested. Beads were washed three more times with 20 mM TEAB and supernatants harvested. Harvested supernatants were vacuum dried before resuspension in 2% acetonitrile and 0.05% trifluoracetic acid for analysis by liquid chromatography tandem mass spectrometry (LC-MS/MS). Samples were acquired in data-dependent acquisition mode and proteins identified using FragPipe based on the mouse proteome database from UniProt (UP00000589) with decoys added by FragPipe. Proteins were quantified using Label-free Quantification. Following analysis, proteins were log_2_ transformed and filtered using Perseus software such that proteins with at least three valid values in either the CpG-stimulated or unstimulated samples were retained for further analysis. Samples were subjected to a two-tailed Student’s T test with a permutation-based false discovery rate of 5%.

### S-Trap trypsin digest protocol for LC-MS/MS

Samples were supplemented with SDS to a final concentration of 5% (activity-based profiling global proteomics). Total protein (20 μg) from activity-based profiling or global proteomics samples were aliquoted and reduced using 10 mM TCEP for 15 minutes at 55⁰C. Following reduction, samples underwent alkylation using 50 mM iodoacetamide for 45 minutes at room temperature in the dark and were then acidified using phosphoric acid to a final concentration of 2.5% v/v. Bind/wash buffer (100 mM TEAB pH 8 and 90% [v/v] methanol) was added and samples were loaded in S-Trap Micro columns (Protifi). Samples were washed a further three times with bind/wash buffer before addition of 1 μg trypsin protease (ThermoFisher) in 20 μL 50 mM TEAB pH 8. Samples were digested for 15 hours at 37⁰C before three elutions of digested peptides using elution buffer 1 (50 mM TEAB), elution buffer 2 (0.2% formic acid) and elution buffer 3 (50% acetonitrile). Samples were dried by vacuum centrifugation and resupended in 2% acetonitrile and 0.05% trifluoracetic acid at a concentration of 0.5 μg/μL for mass spectrometry analysis.

Samples were acquired in data-dependent acquisition mode and proteins identified using FragPipe using the default label-free quantification method. Following analysis, data was log_2_ transformed in Perseus software. Missing values were imputed by downshifting 1.8 standard deviations from the mean, with a variance of 0.3 standard deviations. Samples were subjected to a two-tailed Student’s T test with a permutation-based false discovery rate of 5%.

### Steptavidin blotting

Samples were collected according to activity-based profiling or co-immunoprecipitation protocol. Samples were heated in SDS sample buffer at 95⁰C for 5 minutes to denature proteins. Blotting was performed on affinity purified or whole cell lysates from equal protein concentrations as determined by BCA assay. Proteins were resolved on a pre-cast Bolt 4-12% Bis-Tris Mini Protein gel run with 1X Bolt MES SDS running buffer (Invitrogen). Proteins were transferred onto PVDF membrane with iBlot 2 at 20 volt for seven minutes (Invitrogen). Blots were blocked in PBS-T containing 3% BSA (Sigma) for two hours at room temperature with agitation. For biotin detection, blots were incubated with streptavidin-HRP in PBS-T 1% BSA for 45 minutes at room temperature with agitation. Chemiluminescence was detected using the ECL select substrate (Amersham) acquired on a ChemiDoc MP imaging system (Bio-Rad).

## Acknowledgements

We thank the Melbourne Mass Spectrometry and Proteomics Facility of The Bio21 Molecular Science and Biotechnology Institute for access to MS instrumentation. Many thanks to Professor Edward Tate of Imperial College London supplying IMP-2373 for use in experiments.

---

**Supplementary Figure 1: Bio-Ub-Prg is specific for active DUBs in the lysate of MutuDCs.**

MutuDCs were stimulated with 0.5 μM CpG for 18 hours at 37⁰C. Cells were lysed and lysate protein was quantified by bichinoic acid assay. 500 μg of lysate protein was incubated with 0.1 μg Bio-Ub-Prg activity-based probe (ABP) for 1 hours at 37⁰C. Additional samples were pre-incubated with N-ethylmaleimide (NEM) for 1 hours prior to ABP addition. Streptavidin-agarose beads were used to pulldown probe-labelled proteins. Streptavidin-agarose beads were subjected to on-bead trypsin digest prior to analysis by mass spectrometry. Only DUBs with activity detected in a sample from both treatment groups are shown. Data from four independent experiments, two-tailed Student’s T test with permutation-based false discovery rate with a significance threshold of p_adj_>0.05 (-log_10_(p_adj_ value)>1.3).

**Supplementary Figure 2: IMP-2373-treated MutuDCs have a unique banding pattern relative to vehicle-treated cells.**

MutuDCs were lysed and lysate protein was quantified by bichinoic acid assay. 500 μg of lysate protein from DMSO- and IMP-2373-treated cells were then subject to CuAAC click chemistry addition of biotin to IMP-2373 labelled-proteins. Streptavidin-agarose beads were used to pulldown biotin-labelled proteins. Beads were then boiled at 95C for 5 minutes to elute bound proteins before loading into well for SDS-PAGE. For whole-cell lysates, equal amounts of protein were loaded into each well. Blot representative of one experiment.

## Notes

### Competing Interest Statement

The authors have declared no competing interest.

### Summary of Updates

Manuscript was edited. Tables, Acknowledgments and Funding updated

### Funding

- National Health and Medical Research Council Ideas Grant: 2000783

