## Supplemental Figure 1 for "Identification of active deubiquitinases in dendritic cells under unstimulated and activated conditions"

**Supplementary Figure 1: Bio-Ub-Prg is specific for active DUBs in the lysate of MutuDCs.**

MutuDCs were stimulated with 0.5 μM CpG for 18 hours at 37°C. Cells were lysed and lysate protein was quantified by bichinoic acid assay. 500 μg of lysate protein was incubated with 0.1 μg Bio-Ub-Prg activity-based probe (ABP) for 1 hours at 37°C. Additional samples were pre-incubated with N-ethylmaleimide (NEM) for 1 hours prior to ABP addition. Streptavidin-agarose beads were used to pulldown probe-labelled proteins. Streptavidin-agarose beads were subjected to on-bead trypsin digest prior to analysis by mass spectrometry. Only DUBs with activity detected in a sample from both treatment groups are shown. Data from four independent experiments, two-tailed Student's T test with permutation-based false discovery rate with a significance threshold of $p_{adj}>0.05$ ($-\log_{10}(p_{adj}$ value$)>1.3$).

**Supplementary Figure 2: IMP-2373-treated MutuDCs have a unique banding pattern relative to vehicle-treated cells.**

MutuDCs were lysed and lysate protein was quantified by bichinoic acid assay. 500 μg of lysate protein from DMSO- and IMP-2373-treated cells were then subject to CuAAC click chemistry addition of biotin to IMP-2373 labelled-proteins. Streptavidin-agarose beads were used to pulldown biotin-labelled proteins. Beads were then boiled at 95C for 5 minutes to elute bound proteins before loading into well for SDS-PAGE. For whole-cell lysates, equal amounts of protein were loaded into each well. Blot representative of one experiment.

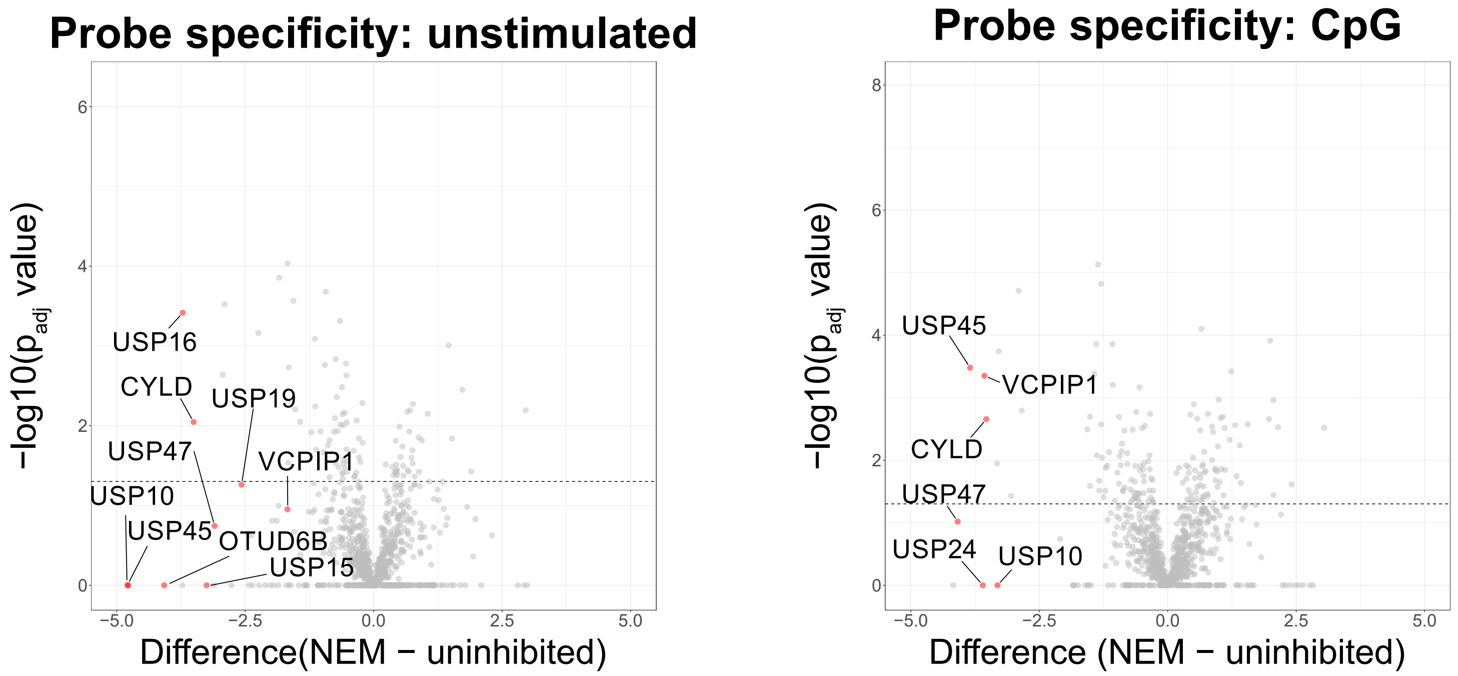


Supplementary Figure 1

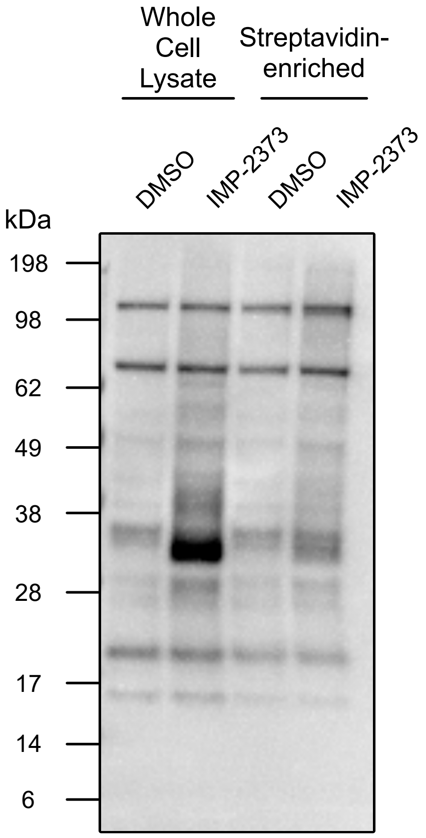


### Supplementary Figure 2
